# Role of MPK4 in pathogen-associated molecular pattern-triggered alternative splicing in Arabidopsis

**DOI:** 10.1101/511980

**Authors:** Jeremie Bazin, Kiruthiga Mariappan, Thomas Blein, Ronny Voelz, Martin Crespi, Heribert Hirt

## Abstract

Alternative splicing (AS) of pre-mRNAs in plants is an important mechanism of gene regulation in environmental stress tolerance but plant signals involved are essentially unknown. Pathogen-associated molecular pattern (PAMP)-triggered immunity (PTI) is mediated by mitogen-activated protein kinases and the majority of PTI defense genes are regulated by MPK3, MPK4 and MPK6. These responses have been mainly analyzed at the transcriptional level, however many splicing factors are direct targets of MAPKs. Here, we studied alternative splicing induced by the PAMP flagellin in Arabidopsis. We identified 506 PAMP-induced differentially alternatively spliced (DAS) genes. Although many DAS genes are targets of nonsense-mediated degradation (NMD), only 19% are potential NMD targets. Importantly, of the 506 PAMP-induced DAS genes, only 89 overlap with the set of 1849 PAMP-induced differentially expressed genes (DEG), indicating that transcriptome analysis does not identify most DAS events. Global DAS analysis of *mpk3*, *mpk4*, and *mpk6* mutants revealed that MPK4 is a key regulator of PAMP-induced differential splicing, regulating AS of a number of splicing factors and immunity-related protein kinases, such as the calcium-dependent protein kinase CPK28, the cysteine-rich receptor like kinases CRK13 and CRK29 or the FLS2 co-receptor SERK4/BKK1. These data suggest that MAP kinase regulation of splicing factors is a key mechanism in PAMP-induced AS regulation of PTI.

**Significance statement:** Alternative pre-mRNA splicing (AS) affects plant responses to environmental stresses. So far, however, the regulation of AS is little understood. Here, we studied AS induced by the pathogen-associated molecular pattern (PAMP) flagellin in Arabidopsis. We identified 506 differentially alternatively spliced (DAS) genes, 89 of which overlap with the 1849 DEG, indicating that the majority of DAS events go undetected by common transcriptome analysis. PAMP-triggered immunity is mediated by mitogen-activated protein kinases. Global DAS analysis of MAPK mutants revealed that MPK4 is a key regulator of AS by affecting splicing factors and a number of important protein kinases involved in immunity. Since PAMP-triggered phosphorylation of several splicing factors is directly mediated by MAPKs, we discovered a key mechanism of AS regulation.

## Introduction

Plants possess pattern recognition receptors that detect conserved pathogen-associated molecular patterns (PAMPs) and initiate PAMP-triggered immunity (PTI) (1). Successful pathogens deliver effectors to the plant apoplast and various intracellular compartments; these effectors suppress PTI and thereby facilitate invasion of the host. As a strategy to counter effectors, plants have evolved intracellular receptors with nucleotide-binding leucine-rich repeat domains that sense effectors and mediate effector-triggered immunity (2).

The bacterial PAMP flg22, a conserved 22-amino-acid peptide derived from *Pseudomonas syringae* flagellin, has provided a powerful tool to decipher PAMP-induced signaling pathways and revealed the complexity of PTI-related mitogen-activated protein kinase (MAPK) cascades. In *Arabidopsis thaliana*, flg22 recognition is mediated by the leucine-rich repeat (LRR) receptor kinase FLAGELLIN SENSITIVE 2 (FLS2). Recognition of flg22 by FLS2 induces an array of defense responses, including the generation of reactive oxygen species (ROS), callose deposition, ethylene production, and reprogramming of host cell genes (1).

Flg22 recognition leads to the activation of two MAPK signaling pathways. One of these MAPK cascades is defined by the mitogen-activated protein kinase kinases (MAPKKs) MKK4 and MKK5, which act redundantly to activate the MAPKs MPK3 and MPK6 (3). The second flg22-activated cascade is defined by the mitogen-activated protein kinase kinase kinase (MAPKKK), MEKK1. This kinase activates the MAPKKs MKK1 and MKK2, which act redundantly on the MAPK MPK4 (4, 5). Flg22 also induces the activity of several other MAPKs, but their functions in plant immunity remain to be clarified (6, 7). Flg22 also transiently activates multiple calcium-dependent protein kinases (CDPKs) in *A. thaliana*. Moreover, four related CDPKs were identified as early transcriptional regulators in PAMP signaling (8). PAMP-induced protein kinase cascades ultimately lead to the regulation of immune response genes to adjust the metabolic and physiological status of the challenged plants. This regulation can occur at the transcriptional or post-transcriptional level, including via AS. Using *mpk3, mpk4*, and *mpk6* mutants, we previously confirmed by microarray-based transcriptomics that PTI defense gene expression is strongly regulated by these three MAPKs (9).

AS of mRNAs is important for stress responses in plants (10, 11). However, very few examples exist how plant signals regulate AS in plant immunity (12–14). Interestingly, proteomic analyses identified a considerable number of splicing-related proteins as major phosphorylation targets in plants (15), and several of these splicing proteins are phosphorylated by MAPKs *in vitro* (16, 17), suggesting a role of MAPKs in AS regulation. Later work confirmed that certain splicing factors are phosphorylated in response to PAMP signaling and carry phosphorylation motifs for CDPKs and MAPKs (18). Recently, we showed that several phosphorylated splicing factors are direct targets of MAPKs (19). For example, MPK4 targets several phosphorylation sites in SCL30 (19). Moreover, *mpk4* mutants are compromised in phosphorylation of the splicing factor SKIP and the RNA helicase DHX8/PRP22, both of which are connected to SCL30 in the phosphorylation network of MPK4 (19).

RNA-sequencing (RNA-seq) allows the assessment of differential expression and AS of genes through quantification of transcripts across a broad dynamic range. Transcript expression levels are inferred based on the number of aligned reads and several methods were specifically developed to evaluate AS events at a genome-wide scale (20). Nevertheless, quantification of transcript isoforms from RNA-seq data remains a substantial challenge. Here, we performed RNA-seq of Arabidopsis treated with flg22. We used standard transcriptome analysis to identify differentially expressed genes (DEGs) and the bioinformatic pipeline AtRTD2 (21) to quantify differential alternative splicing (DAS). These data revealed that 506 genes undergo AS during PTI. However, only 17% of the DAS genes were identified as DEGs. Moreover, DAS genes and DEGs encoded substantially different functional classes of proteins, indicating that current transcriptome analyses miss many regulated transcripts.

Subsequent analysis of *mpk3, mpk4*, and *mpk6* mutants for defects in PAMP-triggered AS revealed that *mpk4* mutant plants were strongly compromised in more than 40% of AS upon PAMP treatment, whereas no major changes in AS transcripts were observed in *mpk3* and *mpk6* mutants under these conditions. Several AS targets involved important rearrangements of protein isoforms of the critical PTI regulators CPK28, CRK29, and SERK4. These results show the genome-wide impact of AS in PTI and the key role MAPKs plays in PAMP-induced AS.

## Results

### Flagellin induces alternative splicing in 506 genes

To identify whether PTI signaling is linked to AS processes in plants, we treated Arabidopsis Col-0 plants for 30 min with flg22 or H_2_O and analyzed the transcripts by RNA-seq from three biological replicates. An average of 46 million 100-nt long paired-end sequencing reads was obtained for each sample. For quality assessment of the RNA-seq data, we analyzed the proportion of read alignments and the genome-wide sequencing coverage. A comparison of the mapped reads with the annotated genes showed that at least 98% of the reads were from exonic, intronic and 5’ or 3’ untranslated regions and only 1% mapped to intergenic regions, respectively (Fig. S1A). Lastly, we assessed the sequencing saturation and found extensive coverage of the chromosomes (Fig. S1B).

We next analyzed the flg22-treated RNA-seq data for DAS events in comparison to mock-treated wild-type plants. We obtained a total of 546 flg22-induced DAS events corresponding to 506 unique DAS genes when considering a p-value cutoff of 0.001 (Fig. 1A), with intron retentions as the most frequent events (85%), followed by 7% and 6% alternative 3’ and 5’ splice sites (3ASS and 5ASS, respectively). Finally, only 6% and 1 single event were obtained for skipped exons (SE) and mutually exclusive exons (MXE), respectively. These results indicate that flg22 induces AS in a considerable number of genes.

**Figure 1:**
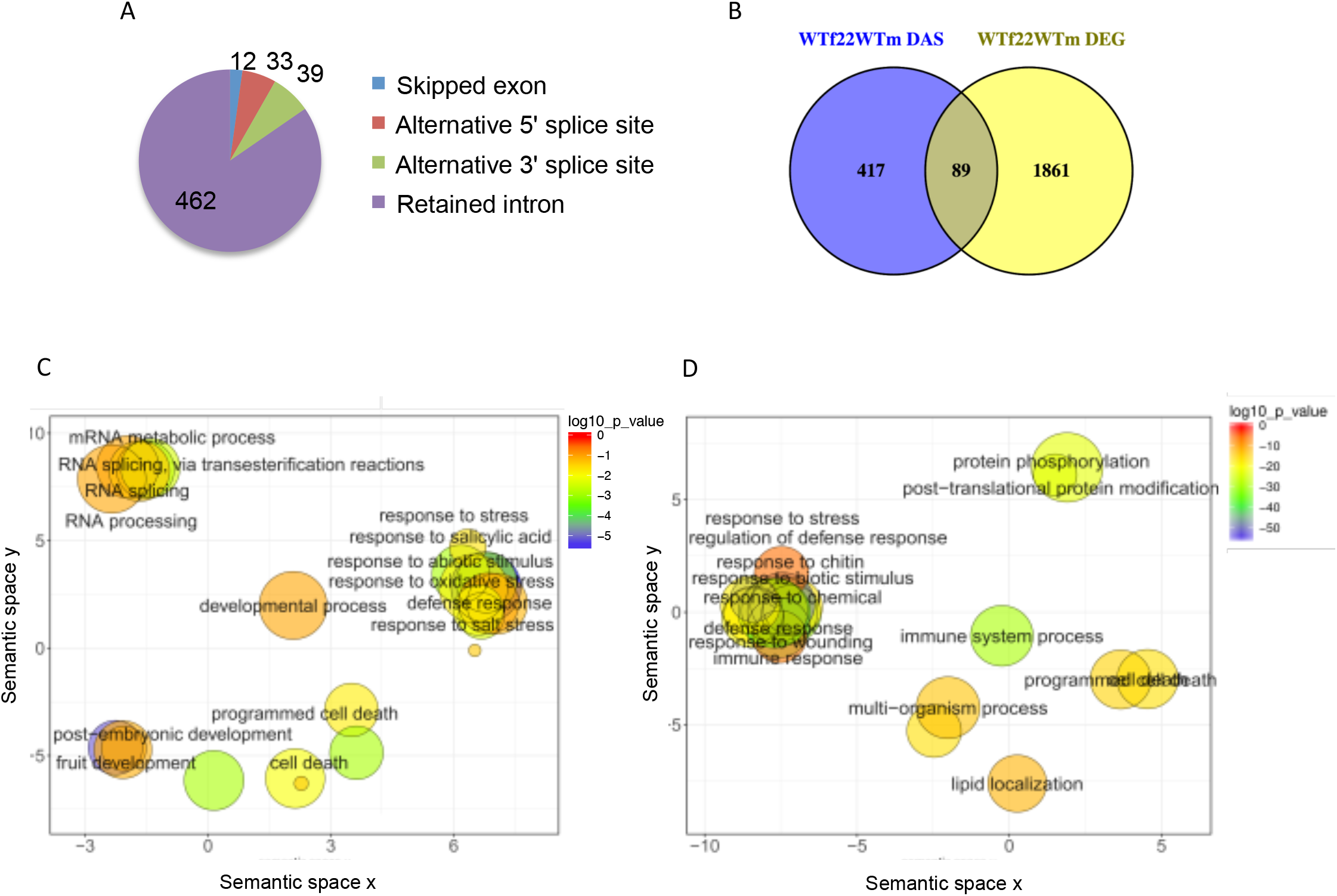
Flagellin induces alternative splicing in 506 genes. **A**, Number of DAS events from each class identified upon flg22 treatment in wild-type plants **B;** Venn comparison plot between differentially spliced (flg22 DAS) with differentially expressed (flg22 DEG) genes in wild-type plants (FDR < 0.01, Fold Change > 2). **C**, REVIGO plots of gene ontology enrichment clusters of differentially spliced, or **D**, differentially expressed genes. Each circle represents a significant GO category but only groups of highest significance are labeled. Related GOs have similar (x, y) coordinates.

When comparing the set of 506 PAMP-induced DAS genes with the set of 1948 DEG (FDR < 0.01, Fold Change > 2) (Fig. 1B; Dataset S1), an overlap of only 89 genes (17%) was obtained. Even when lowering the cutoff value in the set of DEGs to 1.5 (Fig. S2), most DAS genes did not overlap with the DEG data set and hence did not appear as PAMP-regulated genes during classical transcriptome analysis. A comparison of the categories of genes that were differentially spliced (DAS) versus those that were differentially expressed (DEG) showed some overlap in the functional gene ontology (GO) categories of metabolic processes (Fig. 1C, 1D; Fig. S3). Unique enrichment of DAS genes was found in the GO classes of RNA metabolism and development; however, the DAS genes were also classified into functional categories related to plant defense responses and immunity, together with the DEGs (Fig. 1C, 1D).

To reveal the role of RNA metabolism in the set of PAMP-induced DAS genes, we focused on key marker functions in the 506 genes. Indeed, several splicing factors were identified, such as SCL33, SR30, SR45a, RS40, RS41, U2AF65A, RZ1B and RZ1C. Many factors involved in transcriptional regulation, such as RNA-binding proteins, helicases and transcription factors, were found in the set of DAS genes (Dataset S2). The bioinformatic identification of various AS events in a selected number of genes involved in immunity or RNA processing was verified by semi-quantitative RT-PCR and PAGE analysis (Fig. 2).

**Figure 2:**
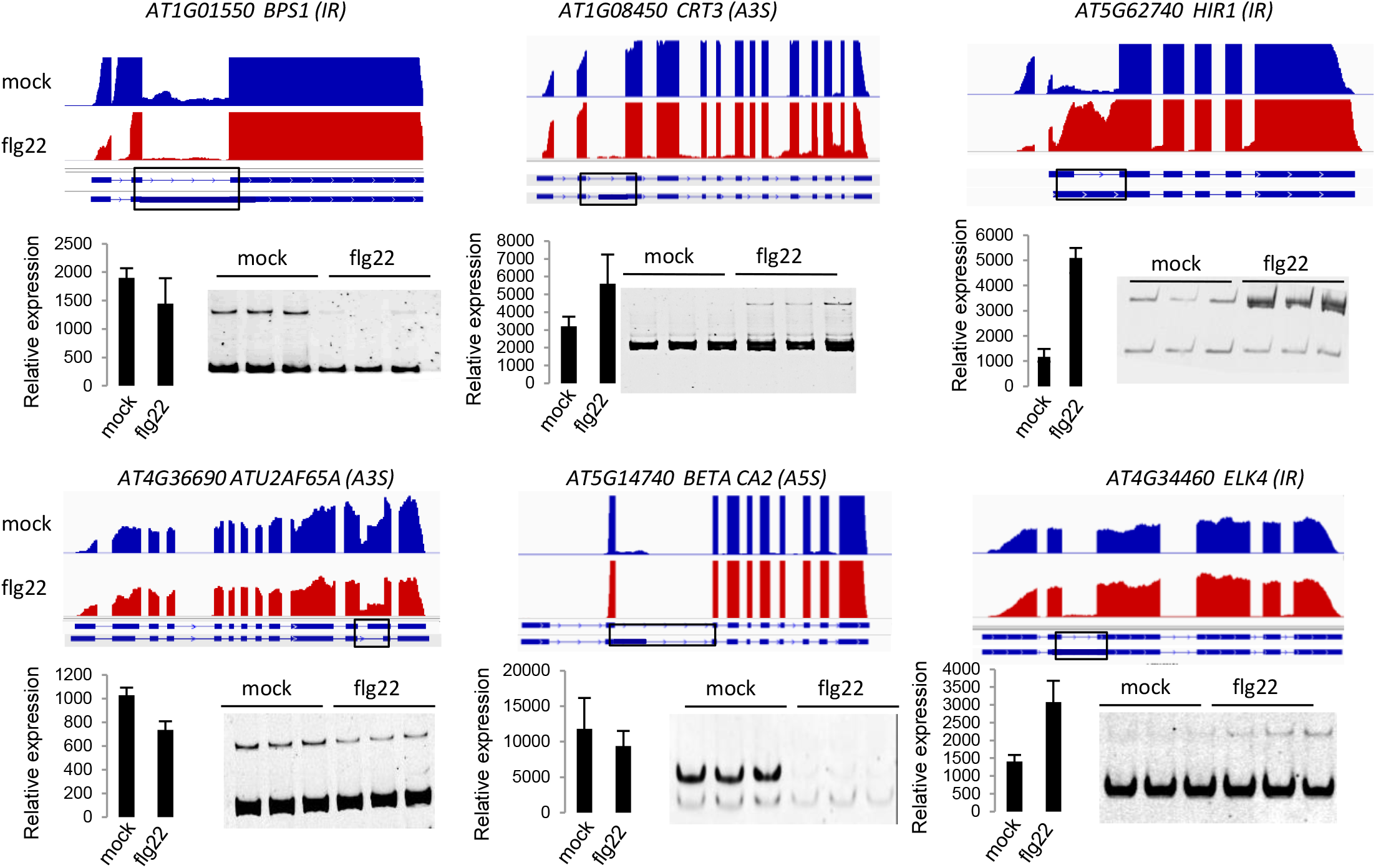
RT-PCR analysis of selected differential splicing events in response to flg22. Normalized RNA-seq coverage data is shown in the upper part of each panel. Scale is the same in each track. SybrGreen stained PAGE gels are shown below. Relative expression for each gene was calculated based on the normalized RNA-seq read counts produced by Deseq2. RT-PCR was performed on three biological replicates.

### MPK4 is a major regulator of PAMP-induced AS

To assess whether the immune-regulated MAPKs, MPK3, MPK4, or MPK6, play a role in regulating PAMP-triggered AS, RNA-seq data were obtained from the three MAPK knock-out mutants before and after flg22 treatment. When we analyzed *mpk3, mpk4*, and *mpk6* in the absence of PAMP treatment, no major changes in AS events were observed when compared to wild type plants (Table 1). However, upon flg22 treatment, we found that PAMP-induced DAS responses were strognly compromised in *mpk4* but not in *mpk3* or *mpk6* (Fig. 3A). Indeed, comparison of the PAMP-induced DAS events observed in *mpk3* and *mpk6* upon flg22 treatment to those in wild-type plants only revealed a total of 12 and 14 DAS genes, respectively (Table 1). Since MPK3 and MPK6 nonetheless show a considerable number of DEGs, we consider the small number of splicing events as minor and conclude that MPK3 and MPK6 are not involved to a significant extent in regulating PAMP-induced AS (Fig. S5). In contrast, comparing *mpk4* plants to wild-type Col-0 in their response to flg22, we identified 419 differential AS events that correspond to 367 unique DAS genes (Table 1, Fig. 3B). Most of the events belong to the class of intron retention (297) followed by 3ASS (30) and 5ASS (28), with only 6 SE (Fig 3B). As analyzed in Venn diagrams, the identity of DAS genes in *mpk4* plants strongly overlapped those seen in flg22-treated wild-type plants (Fig. 3C). Out of the total number of 506 flg22-induced DAS genes in wild type, 39 % (199 genes) were affected in the *mpk4* knock out mutant (Fig. 3C). These results reveal that a major segment of flg22-induced DAS is mediated by MPK4. Importantly, most of the DAS events identified in *mpk4* plants occurred only upon flg22 treatment (Table 1), indicating that the nucleotide-binding and leucine-rich repeat receptor SUMM2, which guards the MPK4 pathway (22), plays no or only a minor role in regulating these events. Consistent with MPK4’s non-immune-related role, 137 DAS genes in *mpk4* mutants were not related to the set of flg22-induced DAS genes in wild-type plants (Fig. 3C).

**Figure 3:**
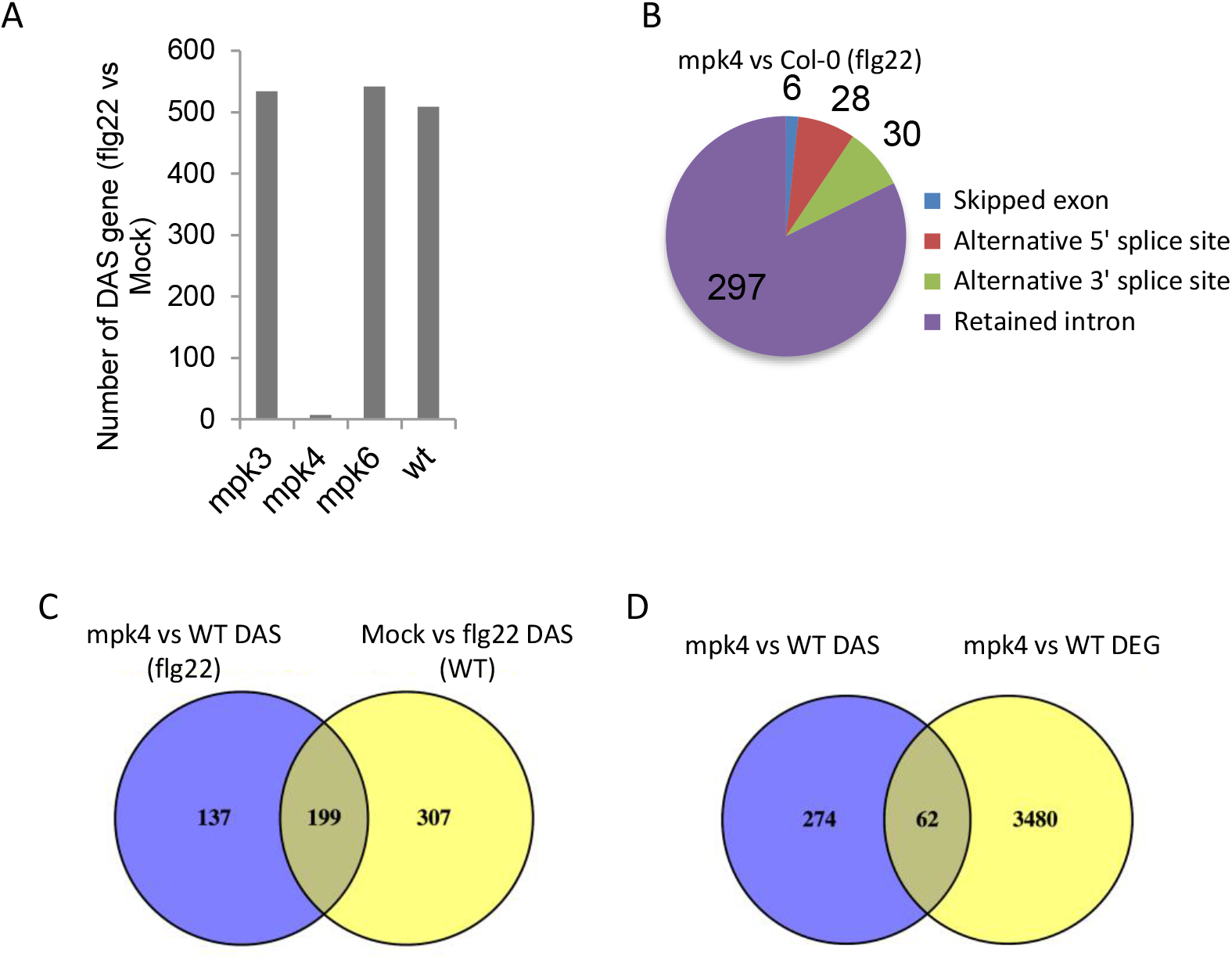
MPK4 is a major regulator of alternative splicing in PTI. **A**, Number of DAS genes in response to flg22 in *mpk3, mpk4, mpk6*, and wild-type (wt, Col-0) plants. **B,** Number of AS events from each class identified in the *mpk4* mutant compared to wild type in the presence of flg22. **C,** Venn comparison plot between DAS genes in *mpk4* in the presence of flg22 with DAS genes in response to flg22 in wild-type plants. **D,** Venn comparison plot between DAS genes and DEG genes in *mpk4* as compared to WT.

**Table 1:** Number and class of DAS events between wild-type (Col) and *mpk3, mpk4*, and *mpk6* upon mock (H_2_O) or flg22 treatment.

| Table 1 : Number of gene with AS events ( Pval cutoff 0.0001 / FDR 0.05) |  |  |  |  |  |
| --- | --- | --- | --- | --- | --- |
| Comparisons vs Col-0 | Skipped exon | Mutually exclusive exon | Alternative 5' splice site | Alternative 3' splice site | Retained intron |
| mpk3 (flg22) | 0 | 0 | 0 | 3 | 8 |
| mpk3 (mock) | 0 | 0 | 2 | 0 | 10 |
| mpk4 (flg22) | 6 | 3 | 28 | 30 | 297 |
| mpk4 (mock) | 3 | 0 | 1 | 13 | 45 |
| mpk6 (flg22) | 7 | 0 | 2 | 0 | 5 |
| mpk6 (mock) | 0 | 0 | 0 | 0 | 10 |

**Table 2:**
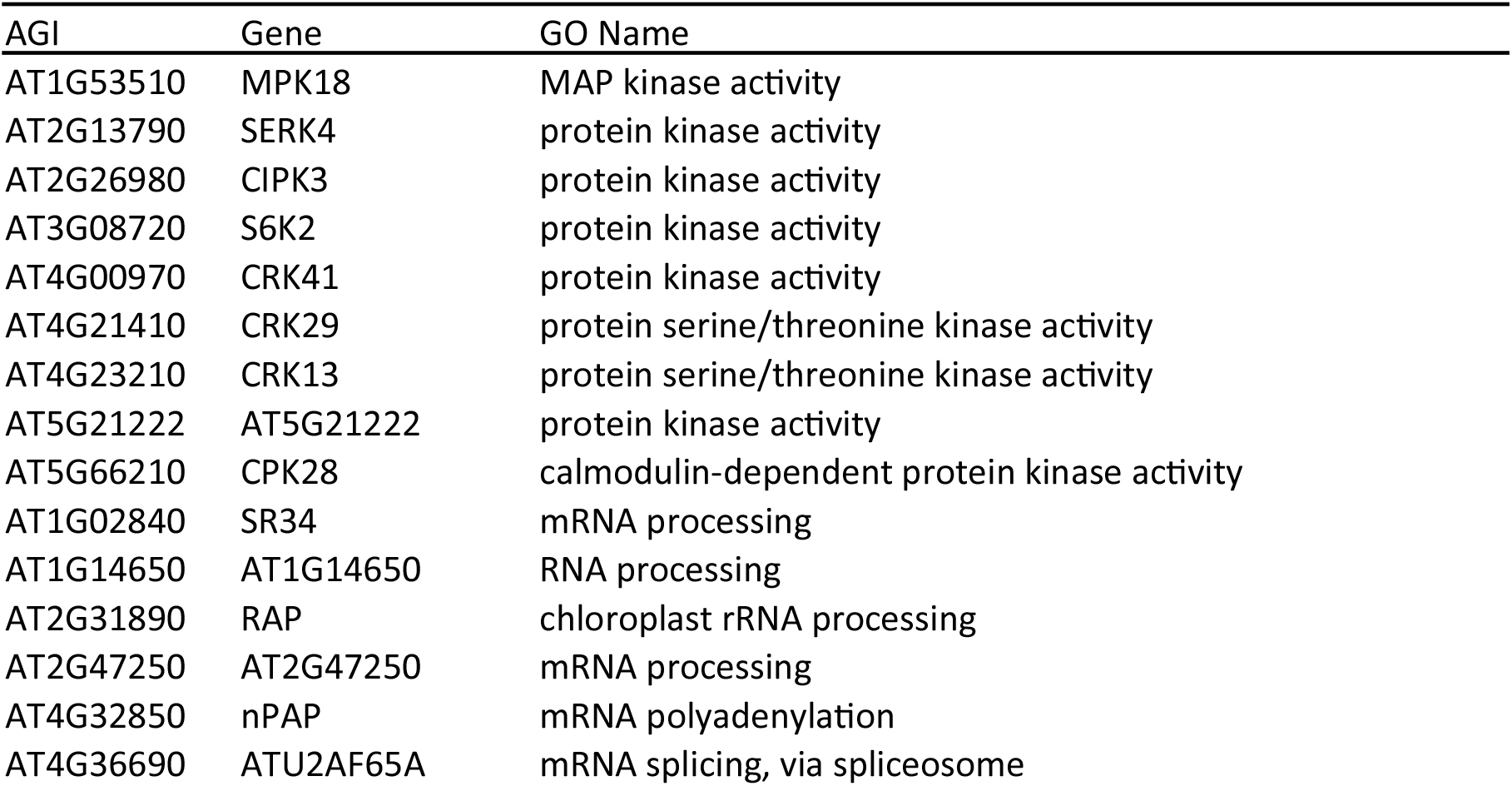
Protein kinases and RNA processing factors with significant isoform-switching events in response to flg22 in wild-type plants. AGI: Gene accession number from the Arabidopsis Genome Initiative.

We next compared the number of DAS genes that overlapped with the DEGs upon treatment of *mpk4* mutants with flg22. As observed in wild-type plants (Fig. 1A), only 18% (62) of DAS genes were found in the DEG dataset when using a two-fold change cutoff value (Fig. 3D), showing that most DAS events are not represented in the set of DEGs. Consistent with the bioinformatics analyses, we found that five out of the six flg22-dependent DAS genes tested by RT-PCR in Figure 2 did not show significant flg22-induced AS in the *mpk4* background (Fig. 4, Fig. S4).

**Figure 4:**
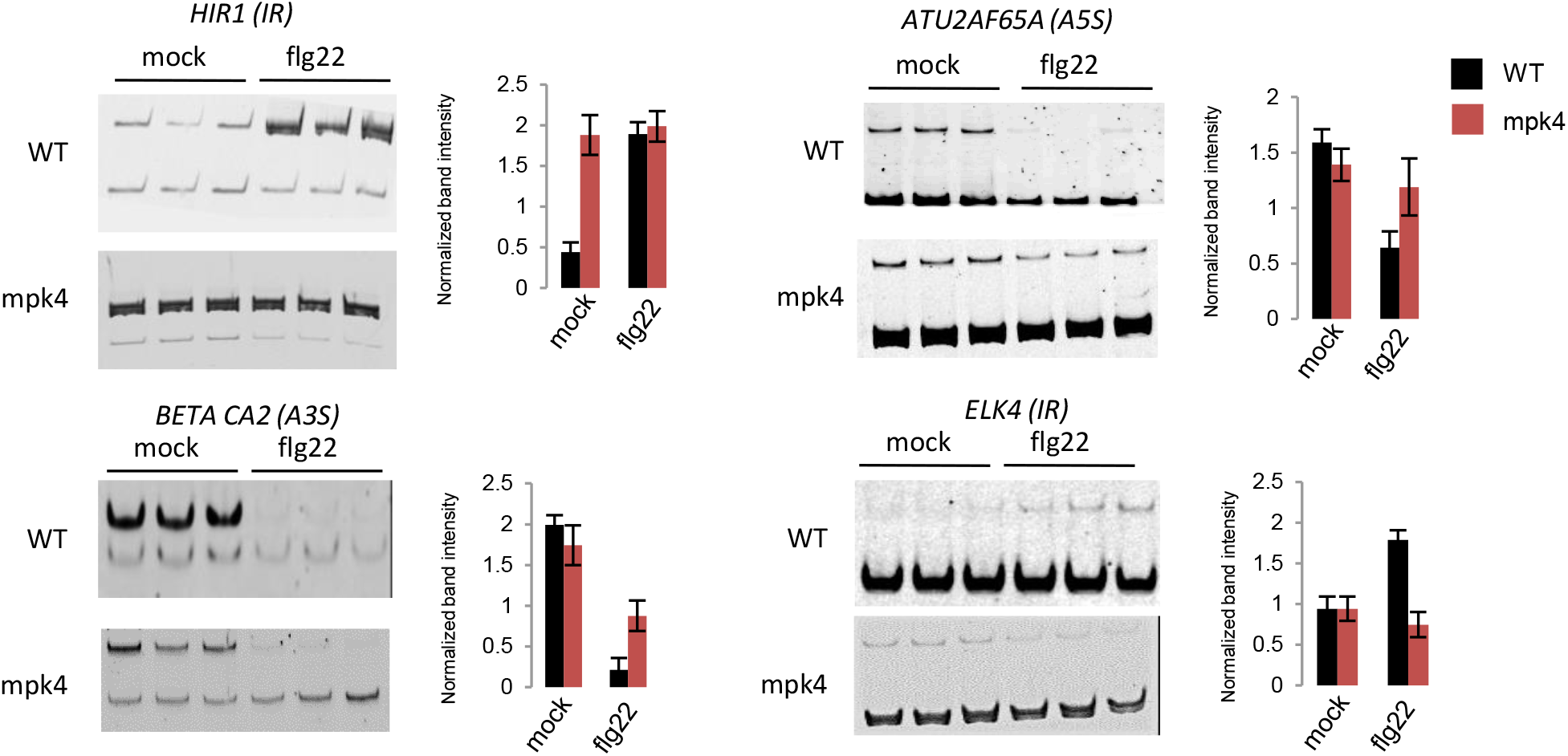
RT-PCR analysis of selected differential splicing events in *mpk4* and WT in mock or flg22-treated plants. RT-PCR was performed on three biological replicates and products were separated on the same gel. Band intensity was normalized against the non-differential isoforms. Significant differences were calculated using a t-test (* p-value < 0.05).

### MPK4 regulates AS of several splicing factor genes

GO enrichment analysis of PAMP-induced DAS targets regulated by MPK4 revealed a strong enrichment for RNA processing and splicing categories (Fig. 5). A set of factors implicated directly in AS exhibited alternative splice site selection in response to flg22, and this AS was dependent on MPK4. Among the DAS splicing factors, several serine/arginine-rich (SR) splicing factors were identified. SR proteins and SR-related proteins are important regulators of constitutive and alternative splicing and other aspects of mRNA metabolism. In the set of PAMP-induced DAS genes that are dependent on MPK4, we identified four nuclear speckle-localized SR proteins At-SCL33, At-SR31, At-RZ1B and At-SR45a, (23–26) (Dataset S2).

**Figure 5:**
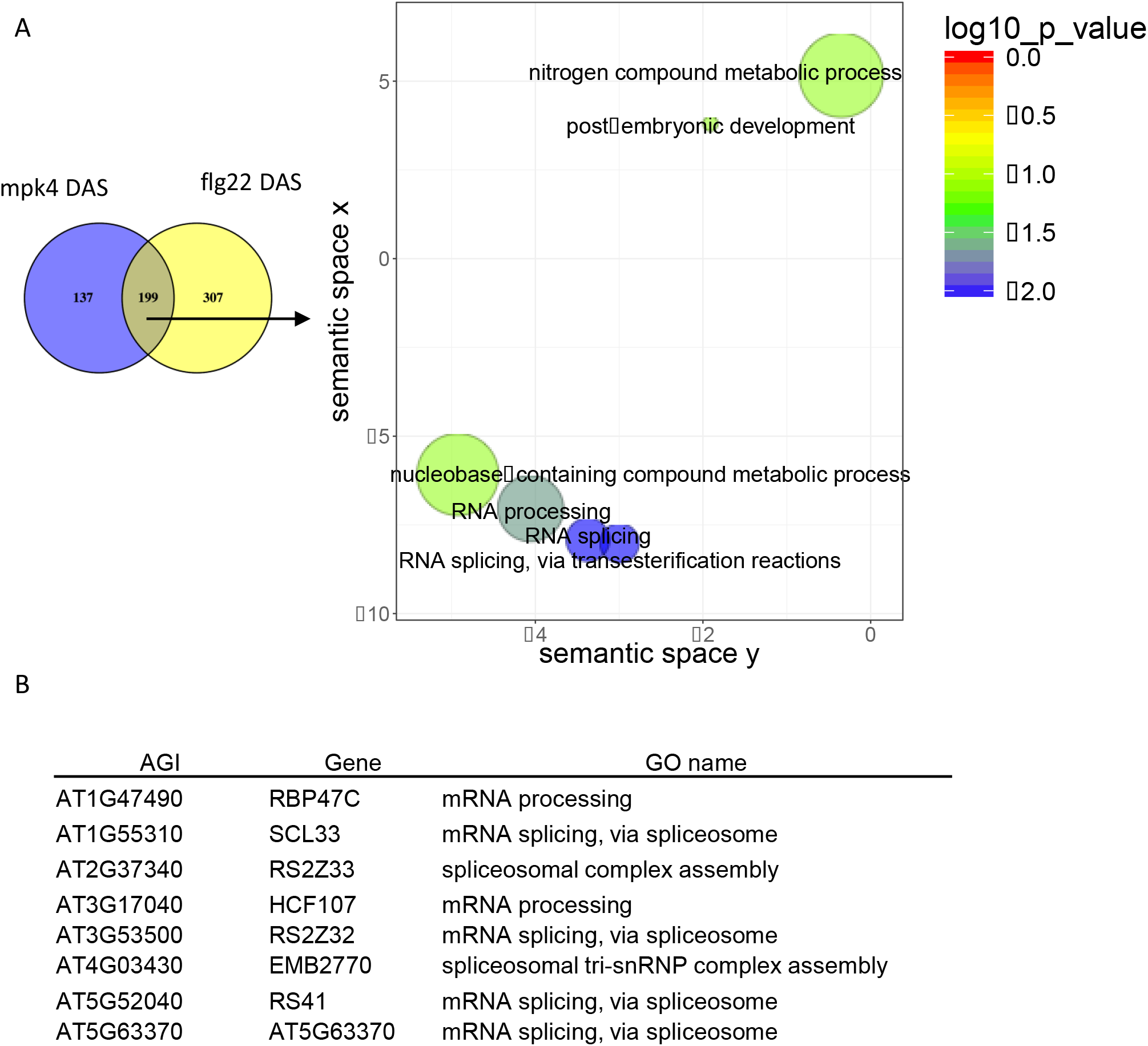
MPK4 regulates alternative processing of several splicing factors. **A,** REVIGO plots of gene ontology enrichment clusters of DAS genes in both *mpk4* compared to wild type and in response to flg22 in wild-type plants. Each circle represents a significant GO category but only groups of highest significance are labeled. Related GOs have similar (x, y) coordinates. **B,** List of DAS genes involved in mRNA processing and splicing.

### Most PAMP-triggered AS transcripts are not targets of nonsense-mediated decay

Many transcripts are AS targets of the nonsense-mediated degradation (NMD) pathway and hence never get translated into protein (27, 28). To assess whether PAMP-triggered AS transcripts are NMD targets, we compared our flg22-induced DAS dataset to that reported by Drechsel et al. (2013) for *upf1 upf3* double mutants, which are defective in the NMD pathway. As shown in Figure 6A, only 108 of the 506 PAMP-induced DAS genes are NMD targets in wild-type Arabidopsis, indicating that 79% of AS transcripts will probably be translated into protein. We concluded that the large majority of PAMP-triggered AS events are not targeted for NMD.

**Figure 6:**
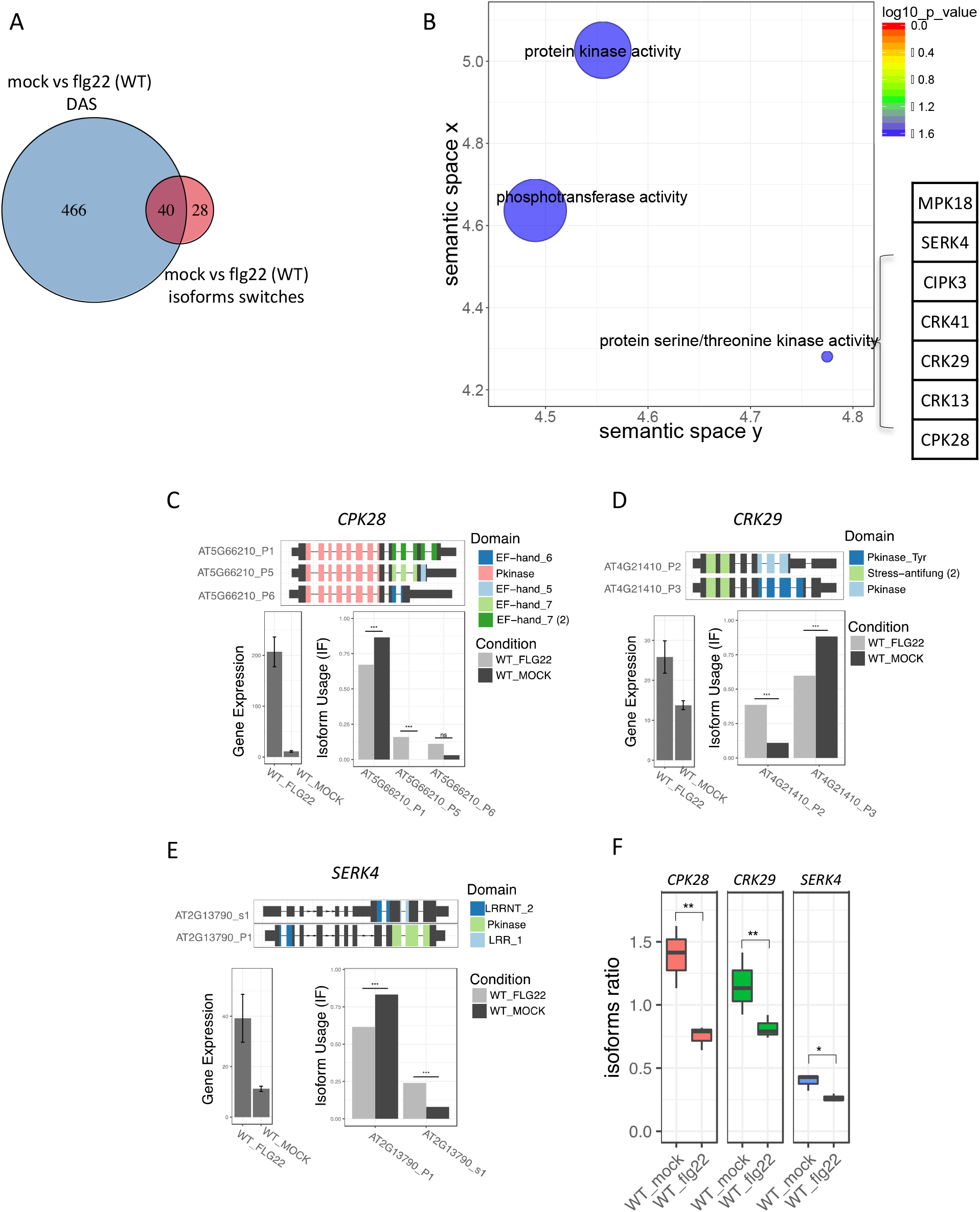
Differentially regulated isoforms are not enriched for NMD targets. **A,** Comparison of PAMP-induced DAS, or **B,** PAMP-induced differentially regulated isoforms, with genes and isoforms up-regulated in *upf1 upf3*, respectively.

### MPK4 regulates PAMP-induced AS of several key stress signaling genes

To analyze the DAS genes affected by MPK4 in relation to pathogen responses, we focused on important immunity-related genes in the list of AS events in *mpk4* (Dataset S2). In the list of PAMP-induced DAS genes that are dependent on MPK4, we identified *RPP4*, which encodes a nucleotide-binding leucine-rich repeat protein with Toll/interleukin-1 receptor domains that confers Arabidopsis resistance to *Peronospora parasitica* (29).

In the MPK4-regulated set of exon-skipping DAS genes, we identified the Ser/Thr protein kinase CIPK3, which associates with a calcineurin B-like calcium sensor, and regulates abscisic acid- and stress-induced gene expression in Arabidopsis (30). The *cipk3* mutants show altered expression of several markers of abscisic acid levels and cold and high salt stress. Another interesting gene showing exon skipping encodes the transcription factor WRKY26 which plays a role in thermotolerance (31). Finally, MPK4 also regulates PAMP-induced AS of *NTH2* gene, encoding a glycosylase-lyase that is involved in oxidative DNA damage repair (32).

In the list of 5ASS genes, *CIPK3* is found again, as well as the MAP kinase gene *MPK17* and the transcription factor gene *WRKY19*, which encodes a MAPKK kinase, but for which no function(s) have yet been attributed.

### PTI is associated with alternative transcript usage of protein kinase isoforms

Functional consequences of AS rely on changes in the use of different isoforms and on the functional differences between the proteins encoded by alternative isoforms. Therefore, we used a combination of recent methods allowing isoform quantification from RNA-seq read pseudo-counts, statistical identification of changes in isoform usage ratio (isoform switching events), and automated prediction of putative functional differences between alternative isoforms. When we used this approach on flg22-treated wild-type plants, we identified 68 genes showing significant isoform-switching events (Fig. 7A; Dataset S3), 48 of which were also identified as DAS by the MATS software. Strikingly, GO analysis of these 68 genes revealed a unique enrichment for protein kinases, some of which are involved in PTI (Fig. 7B). AS of many of these genes is predicted to change the kinase domains of the encoded proteins. For instance, as shown in Figure 7C, we found isoform switching in the mRNA region encoding the kinase domain of CPK28, a calcium-dependent protein kinase that attenuates PTI by phosphorylating BIK1, which is a cytoplasmic protein kinase-mediating multiple pattern recognition receptor (35). Similarly, two cysteine-rich receptor-like kinase (CRK) genes, *CRK13* and *CRK29*, (Fig. 7D) displayed a conserved intron retention events in their kinase domains, leading to increased usage of isoforms containing both serine/threonine and tyrosine kinase domains. Interestingly, these two CRKs differentially influence the sensitivity to infection by pathogenic *Pseudomonas syringae* strains (36). Finally, *SOMATIC EMBRYOGENESIS-RELATED KINASE4* (*SERK4*/*BKK1*), which encodes a FLS2 co-receptor together with its closest homolog BAK1 (37–39), was transcriptionally induced by flg22 but showed a decrease in the isoform predicted to contain a functional kinase domain (Fig. 7E). Isoform-specific RT-qPCR analysis of *CPK28, CRK29*, and *SERK4* confirmed the bioinformatically detected changes (Fig 7F). To assess whether the PAMP-induced AS isoforms are NMD targets, we comparing the PAMP-induced set of isoform-switching events with those targeted by the NMD pathway (Fig. 6B). However, only 10% of the AS isoforms were found to be targets of NMD, indicating that PAMP-induced protein kinase isoforms will likely contribute to PTI.

**Figure 7:**
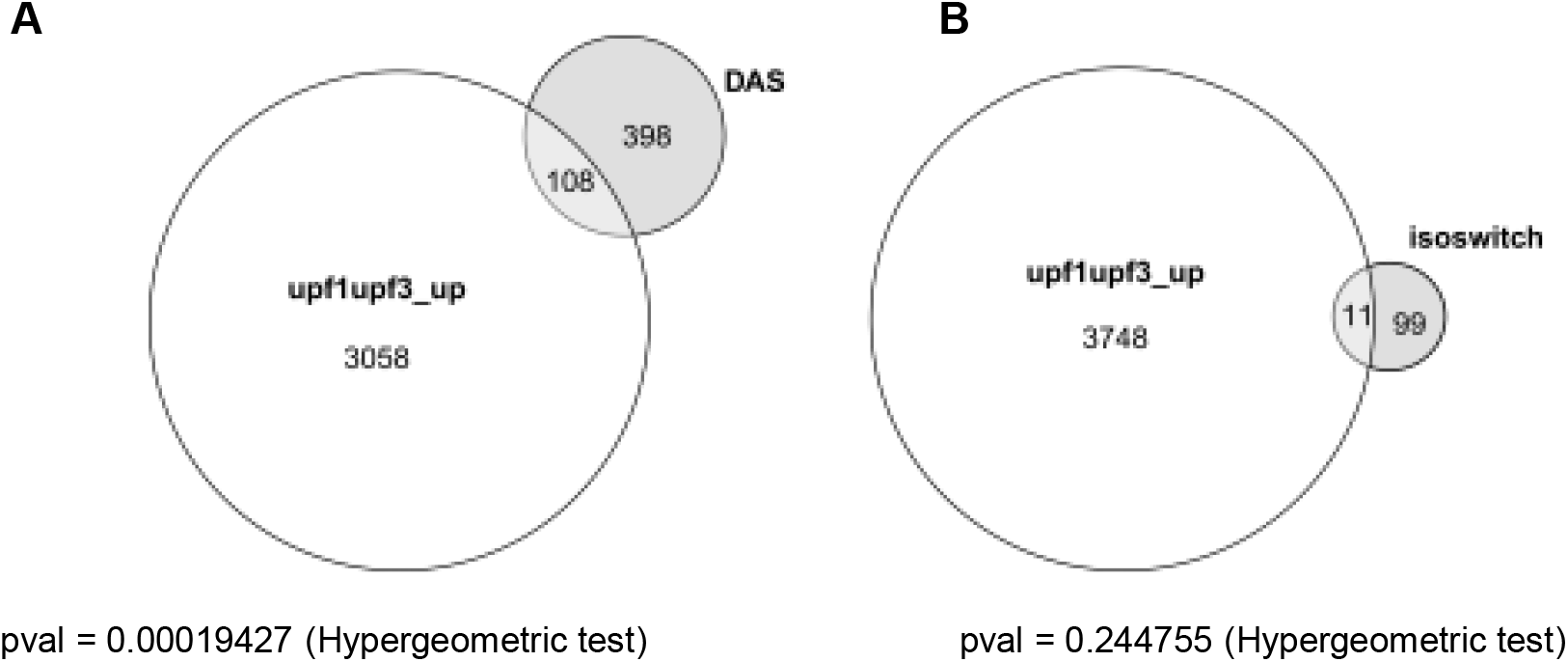
PTI is associated with alternative usage of transcript isoforms of protein kinase genes involved in PTI. **A,** Venn comparison plot between DAS genes compared to genes with isoforms switching events in response to flg22 in WT. **B,** REVIGO plots of gene ontology enrichment clusters of DAS genes both in *mpk4* compared to wild type and in response to flg22 in wild-type plants. Each circle represents a significant GO category but only groups of highest significance are labeled. Related GOs have similar (x, y) coordinates. **C,** Results of isoform switch analysis showing the gene diagram of differential isoforms with the predicted PFAM domain, change in gene expression and isoform usage in response to flg22 for CPK28, **D,** CRK29, and **E,** SERK4. **F,** Isoforms specific RT-qPCR analysis showing the ratio of isoform abundances in response to mock (H2O) or flg22 for CPK28, CRK29 and SERK4. RT-qPCR was performed on three biological replicates. Significant differences were estimated with a student t-test (** p-value < 0.01; * p-value < 0.05).

## Discussion

In higher plants, AS plays a key role in gene expression as shown by the fact that 60–70% of intron-containing genes undergo alternative processing (40, 41). AS is important in normal growth, development, as well as in abiotic and biotic stress responses in Arabidopsis (10–14), but little is known how plant signals trigger AS events and their impact in rapid changes in gene regulation.

### PAMP-triggered AS of defense genes

PAMP-induced protein kinase cascades regulate the expression of immune response genes to adjust the metabolism and physiological status of plants. We compared the set of 1849 PAMP-induced DEG with that of 506 DAS genes. Only 89 of the 506 DAS genes were included in the set of DEGs. Moreover, PAMP-induced DAS and DEG encoded substantially different functional classes of proteins. DAS genes showed unique enrichment for roles in RNA metabolism and transcription, whereas DEGs showed unique functions in responses to various stimuli, signaling and defense. To reveal the role of RNA metabolism functions in the PAMP-induced DAS set of genes, we searched for key markers in the 506 DAS genes. Although most transcripts showed intron retention, several splicing factors were identified as alternatively spliced isoforms in 5ASS, 3ASS, and SE transcripts. Among these PAMP-induced DAS genes, we found the splicing factors RS40, RS41, SCL33, SR30, SR45A, U2AF65A, RZ1B, and RZ1C.

Apart from genes with a direct role in RNA metabolism, we identified several PAMP-induced AS transcripts that encode factors involved in defense and signaling. Leading the list of PAMP-induced AS genes are receptor-like kinases, such as the cysteine-rich receptor-like kinases *CRK13* and *CRK29*, or the FLS2 co-receptor BIK1/SERK4. RLKs and immune receptors in general seem to be a preferred target of immunity-related AS events in plants (43). Among these, two AS isoforms of the TIR-NB LRR receptor N are important for TMV resistance (44) and AS also regulates the barley and rice CC-NB-LRRs Mla13 and Pi-ta, respectively. In barley, powdery mildew induced defense is linked to the AS ratio of five different Mla13 splice isoforms (45), whereby in rice, eleven out of the twelve Pi-ta splice variants encode different proteins (46). In Arabidopsis, RPS4 presents a similar complicated case of six splice isoforms (47). In many cases, the expression of different isoforms of these receptor-like proteins plays a critical role in defense activation via shaping their intramolecular and intermolecular interactions with themselves and other proteins (43). However, so far, the underlying molecular mechanisms which control AS of these genes is poorly understood, but genetics provides some interesting leads. For example, mutation of the SR-type splicing regulator MOS12 affects AS of the Arabidopsis NB-LRRs SNC1 and RPS4 (47). Besides a direct involvement of splicing regulators, AS could also be affected by the epigenetic state of the respective genes, their transcription rates and other post-transcriptional events.

Many of the PAMP-induced defense responses are mediated by the activation of transcription factors and several different classes of transcription factors have been found to show PAMP-induced AS. Among these, the WRKY family plays a key role in abiotic and biotic stresses (48). Here, we identified *WRKY26* as a target of PAMP-induced AS. WRKY26 functions with WRKY33, which is a key regulator of PTI (1) in response to heat stress (31). Our results support the notion that AS of these WRKY transcription factors likely contribute to immunity regulation, a notion that has been proven for the rice WRKYs OsWRKY62 and OsWRKY76 which undergo AS and thereby alter the DNA binding properties and functions in plant defense (49).

Most plant defenses against microbes are based on PAMP-induced production of ROS (50) and a number of enzymes in ROS production contribute directly to disease resistance (51). Interestingly, besides *CATALASE 3*, a key enzyme in ROS detoxification and a target of virus-induced necrosis (34), we also identified *FQR1* in the set of PAMP-induced DAS genes. FQR1 is a cytosolic quinone reductase that protects Arabidopsis from necrotrophic fungi (33). Knock-out lines of *fqr1* displayed significantly slower development of lesions of *Botrytis cinerea* and *Sclerotinia sclerotium* in comparison to the wild type. Consistent with a role in disease resistance, *fqr1* mutants displayed increased ROS accumulation and defense gene expression and overexpression of *FQR1* resulted in hypersensitivity to pathogens (33).

### MPK4 is a major regulator of alternative splicing in PAMP-triggered immunity

The immune MAPKs MPK3, MPK4, and MPK6, are key players of PAMP-triggered gene expression (1). We therefore tested whether any of the three mutant MAPKs might also affect these rapid PAMP-induced AS events. In contrast to MPK3 and MPK6, MPK4 was found to be a major regulator of the AS regulation in response to flagellin. Importantly, about 40% of the AS events that were induced by flg22 in wild-type plants seem to be regulated by MPK4, revealing that MPK4 is a key regulator in AS during pathogen defense. Mutants of MPK4 were compromised in AS events of all categories with 85% intron retention events, followed by 8% 3ASS and 5ASS, 4% SE and 2% MXE.

A number of splicing-related proteins are PAMP-induced phosphorylation targets of MAPKs *in vitro* and *in vivo* (15, 17–19). Among these, we identified SCL30 as a target of MPK4. *scl30* mutants are compromised in phosphorylation of the splicing factor SKIP and the RNA helicase DHX8/PRP22, both of which are connected to SCL30 in the phosphorylation network of MPK4 (19). We also found that flg22-induced AS of the SR protein *At-RS31* is dependent on MPK4. AS of *At-RS31* was also shown to be modulated by dark/light transition through a chloroplast retrograde signaling pathway (24), suggesting that it is a central AS event in several signaling pathways. Moreover, *mpk4* mutants were also compromised in flg22-induced splicing of *At-RZ1B*, which interacts with a spectrum of SR proteins (25). RZ-1B localizes to nuclear speckles and interacts with several SR proteins and loss-of-function of *At-RZ1B* is accompanied by defective splicing of many genes and global perturbation of gene expression (21). We also found the putative splicing factor At-SR45a, which shows differential 3′ splice site selection in the *mpk4* mutant upon flg22 treatment. SR45a can interact with U1-70K, U2AF (35)b, SR45, At-SCL28, and PRP38-like proteins and undergoes AS itself. U1-70K and U2AF (35)b are splicing factors that play a critical role in the initial definition of 5′ and 3′ splice sites and in the early stages of spliceosome assembly. SCL28 and PRP38-like protein are homologs of the splicing factors essential for cleavage of the 5’ splice site (Tanabe et al., 2009). The N-terminal extension in the splice form of the SR45A-1a protein inhibits interaction with these splicing factors, suggesting that SR45A helps to form the bridge between 5′ and 3′ splice sites in the spliceosome assembly and the efficiency of spliceosome formation is affected by the expression ratio of SR45a-1a and SR45a-2 (26).

As MPK4 regulates the splicing of several AS regulators, this consequently adds to the plasticity and complexity of the AS network triggered during pathogen defense. Primary effects of PAMP-triggered signaling by MPK4 on these AS regulators may lead to secondary AS targets, as has been shown for rapid regulation of RS33 by the chloroplast retrograde signaling (24) and for certain auxin targets (42). This may hint to the existence of AS feedback control mechanisms on these AS-controlled regulators similar to those known for transcriptional circuits.

## Conclusions

Our data suggest that PAMP-induced AS could be directly regulated by MAPK phosphorylation of splicing factors that may generate important isoforms of key pathogen receptors, signaling components and enzymes controlling plant defense. This work warrants further investigation of AS regulation in plant defense and signaling.

## Materials and Methods

### Plant materials and treatments

*Arabidopsis thaliana* ecotype Col-0 was used as wild type. The MAPK mutants were *mpk4-2* (SALK_056245), *mpk3* (SALK_151594), and *mpk6-2* (SALK_073907). Seeds were surface-sterilised and stratified for 2 d at 4 °C. Seedlings were then grown for 13 d in a culture chamber at 22 °C with a 16 h light photoperiod, on MS plates (0.5 × Murashige Skoog Basal Salts (Sigma #M6899), 1% sucrose, 0.5% agar, 0.5 % MES, pH 5.7). Twenty-four h before treatment, liquid MS (same media without agar) was added to the MS plates to facilitate the transfer of seedlings to liquid MS. Seedlings were treated with deionized water (mock) or with a final concentration of 1 μM flg22 for 30 min and then frozen in liquid nitrogen. In the case of the *mpk4* single mutant, the *mpk4-2* mutation was segregating. These seedlings were thus first grown vertically in MS plates with 1% agar for 7 d to isolate *mpk4-/-* seedlings based on their root phenotype (thickening and shortening of the primary root [82]). Selected seedlings were then grown for another 6 d at 22 °C with a 16 h light photoperiod, on MS plates before transfer to liquid MS and treatments as described above for the other lines.

### RNA extraction, library construction, and sequencing

Three independent biological replicates were produced. For each biological repetition and each timepoint, 14-day-old seedlings grown in long day conditions were collected and RNA samples were obtained by pooling more than 50 plants. Total RNA was extracted with NucleoSpin RNA Plant (MACHEREY-NAGEL), according to the manufacturer’s instructions. First-strand cDNA was synthesised from 5 μg of total RNAs using the SuperScript First-Strand Synthesis System for RT-PCR (Life Technologies), according to the manufacturer’s instructions. The cDNA stock was diluted to a final concentration of 25 ng/μl. Subsequently, 500 nM of each primer was applied and mixed with LightCycler 480 Sybr Green I Master mix (Roche Applied Science) for quantitative PCR analysis, according to the manufacturer’s instructions. Products were amplified and fluorescent signals acquired with a LightCycler 480 detection system. The specificity of amplification products was determined by melting curves. *GADPH* was used as internal control for signal normalization. Exor4 relative quantification software (Roche Applied Science) automatically calculated the relative expression level of the selected genes with algorithms based on the ΔΔCt method. Data were used from duplicates of at least three biological replicates. Sequencing was performed on each library to generate 101-bp paired-end reads on the Illumina HiSeq4000 Genome Analyzer platform. Read quality was checked by the use of FastQC (52) and low quality reads were trimmed by the use of Trimmomatic version 0.32 (http://www.usadellab.org/cms/?page=trimmomatic) with the following parameters: minimum length of 36 bp; mean Phred quality score higher than 30; leading and trailing base removal with base quality below 3; and sliding window of 4:15. After pre-processing the Illumina reads, the transcript structures were reconstructed by the use of a series of programs, namely, TopHat (ver. 2.1.1; http://tophat.cbcb.umd.edu/) for aligning with the genome, and Cufflinks (ver. 2.2.1; http://cufflinks.cbcb.umd.edu/) for gene structure predictions. For TopHat, the Reference-*Arabidopsis thaliana* (TAIR10) genome (https://www.arabidopsis.org) was used as the reference sequences with the maximum number of mismatches set to two. Raw RNA-seq data files are available at the European Nucleotide Archive (ENA) under the accession number PRJNA379910 (https://www.ebi.ac.uk/ena/) with the following accession numbers SRR5363192 (MP6mock_3), SRR5363193 (MPK6mock_2), SRR5363194 (MPK6mock_1), SRR5363195 (MPK6flg22_3), SRR5363196 (MPK6flg22_2), SRR5363197 (MPK6flg22_1), SRR5363198 (MPK3mock_3), SRR5363199 (MPK3mock_2), SRR5363200 (MPK3mock_1), SRR5363201 (MPK3flg22_3), SRR5363202 (MPK3flg22_2), SRR5363203 (MPK3flg22_1), SRR5363204 (MPK4mock_3), SRR5363205 (MPK4mock_P2), SRR5363206 (MPK4mock_1), SRR5363207 (MPK4flg2_3), SRR5363208 (MPK4flg22_2), SRR5363209 (MPK4flg22_1), SRR5363210 (WTmock_3), SRR5363211 (WTmock_2), SRR5363212 (WTmock_1), SRR5363213 (WTflg22_3), SRR5363214 (WTflg22_2) and SRR5363215 (WTflg22_1).

### Bioinformatic analysis of RNA-seq data

Reads were quality-checked using FASTQC (52) and trimmed using Trimmomatic V0.36 (53) to remove low-quality reads/bps. Trimmed reads were then aligned to the Arabidopsis reference genome (TAIR10) (54) with AtRTD2 database (21, 55) using Tophat v2.1.1 (56-58). Differential expression from the aligned reads was calculated using the cufflinks V2.2.1 package (57). Genes with a two-fold change and a p-value cutoff of 0.05 were considered to be differentially expressed. Genes that were commonly DE and DAS are identified using Venny 2.1.0 (59).

Wild-type, *upf1, upf3* and *upf1 upf3* RNA-seq datasets from (28) were downloaded from the ENA (https://www.ebi.ac.uk/ena) under the following accession numbers SRR584115 (WT-1), SRR584121 (WT-2), SRR584116 (upf1-1), SRR584122 (*upf1-2*), SRR584118 (upf3-1), SRR584124 (upf3-2), SRR584117 (*upf1 upf3-1*) and SRR584123 (*upf1 upf3-2*). Transcript isoform abundance was quantified with pseudo-alignment read count with *kallisto* (60), on all isoforms of the AtRTD2 database (21) and the Araport11 annotation database (https://www.araport.org/data/araport11). Differential expression analysis was performed, both at the transcript or gene level, with DEseq2 with Bonferroni correction of the p-value. Transcripts or genes significantly up-regulated in *upf* mutants compared to Col-0 (p adjusted < 0.01, logFC > 1) were considered as potential NMD targets. The significance of the overlap of NMD targets among DAS genes or isoforms was calculated with a hypergeometric test.

### Alternative splicing analysis

Alternatively spliced genes for wild type and MPK treated/untreated samples were identified using rMATS (61). Only isoforms with a false discovery rate (FDR) cutoff ≤ 0.05 were considered to be significant. For isoform switching identification, transcript isoform abundance was quantified with pseudo-alignment read count with *kallisto* (60) on all isoforms of the AtRTD2 database (21). Then the IsoformSwitchAnalyzeR package was used to detect significant changes in isoform usage. Only significant switches (p adj < 0.1) were kept for further analyses (62).

### AS events validation by RT-PCR

Total RNA was treated with DNAseI (Thermo Fisher Scientific) according to the manufacturer’s instruction and 1 μg of DNA-free RNA was reversed transcribed with an oligo (dT) primer using the Maxima H Minus Reverse Transcriptase (Thermo). cDNA was amplified with primers spanning the splicing events predicted by rMATS, separated on a 6% PAGE gel, stained for 5 min in SybrGold (Thermo) and imaged using the ChemiDoc XRS system (BioRad). Band pixel density was calculated using ImageJ on three biological replicates. The relative band intensity was calculated as the ratio of the AS RT-PCR product versus the constitutive spliced junction product for each replicate. For isoform-specific RT-qPCR, cDNA was produced as described above and qPCR was performed using primers matching non-overlapping regions between isoforms. Significant differences were calculated using Student’s t-test on three biological replicates.

---

**Fig. S1:**
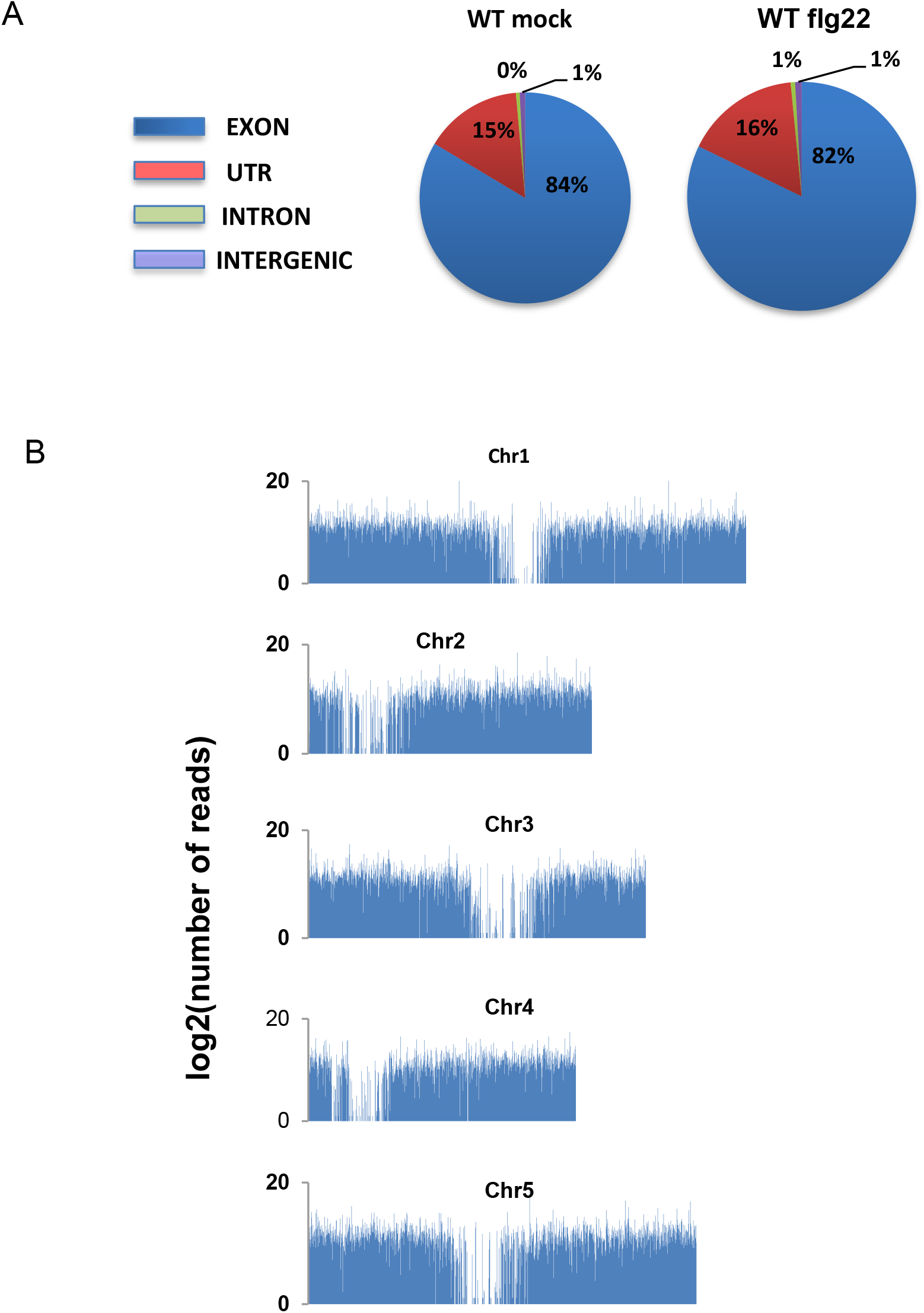
**A,** Percentage of mapped reads to exonic, intronic, 5’ or 3’ untranslated regions (UTR) and intergenic regions in WT upon mock (H_2_O) or flg22 treatment. **B,** Sequencing coverage of chromosomes.

**Fig. S2:**
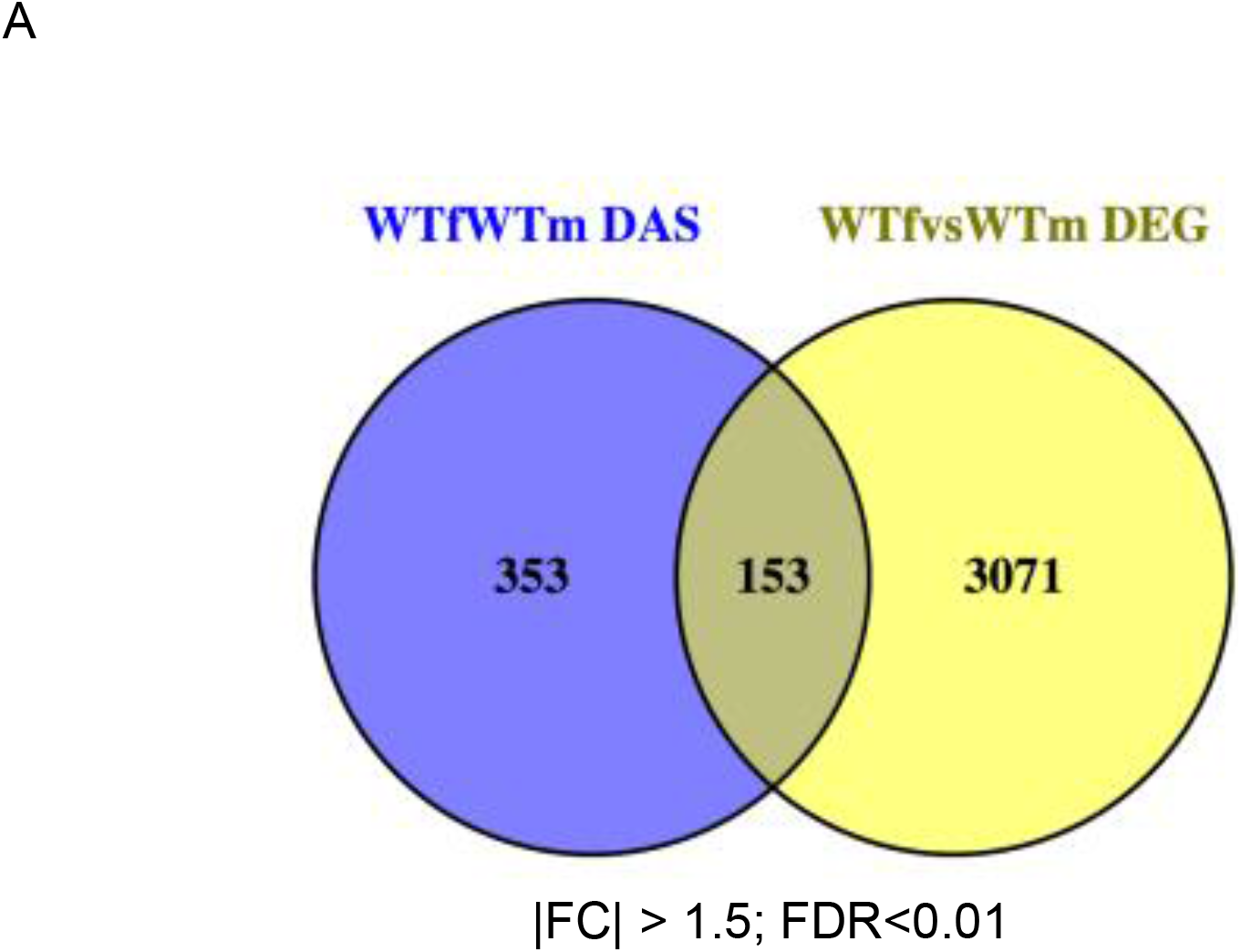
Venn comparison plot between differentially expressed (flg22 DEG) and differentially spliced (flg22 DAS) genes in wild-type plants (FDR < 0.01, Fold Change > 1.5).

**Fig. S3:**
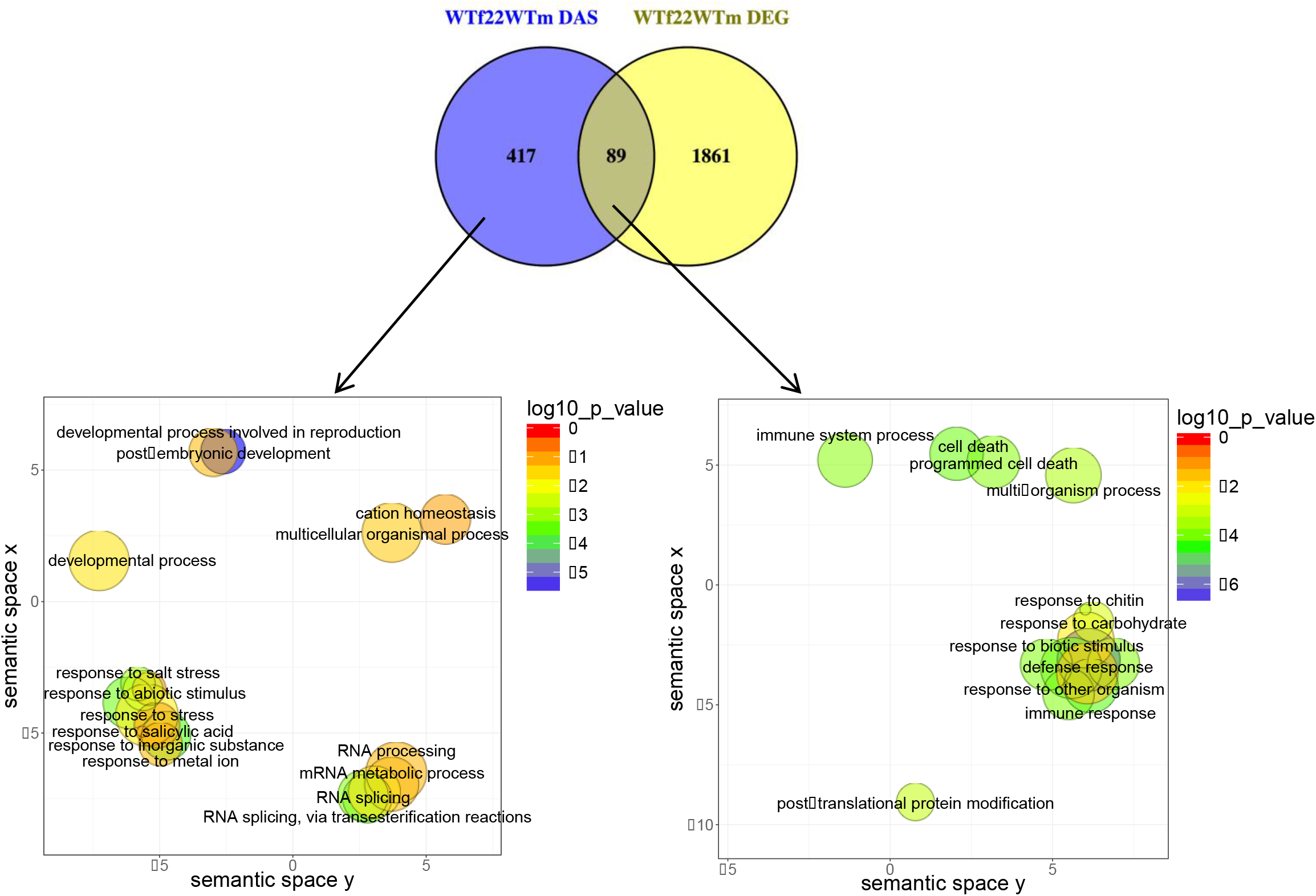
Venn comparison plot between differentially spliced (flg22 DAS) with differentially expressed (flg22 DEG) genes in wild-type plants (FDR < 0.01, Fold Change > 2), showing REVIGO plots of gene ontology enrichment clusters of flg22 DAS-specific and flg22 DAS genes that overlap with flg22 DEG. Each circle represents a significant GO category but only groups of highest significance are labeled. Related GOs have similar (x, y) coordinates.

**Figure S4:**
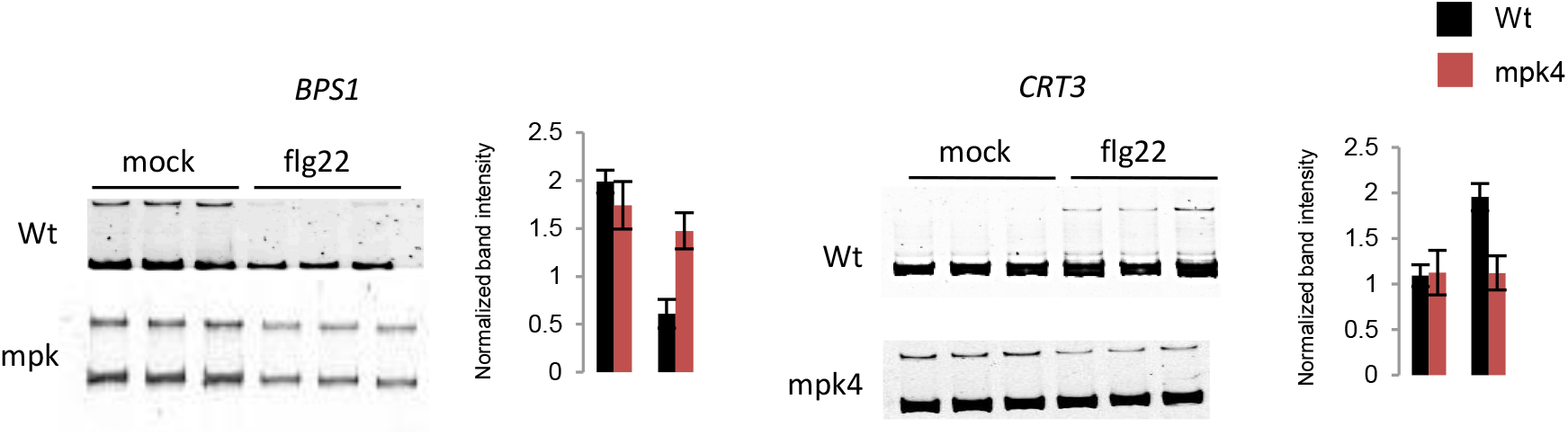
RT-PCR analysis of selected differential splicing events in *mpk4* and WT in mock or flg22-treated plants. RT-PCR was performed on three biological replicates and products were separated on the same gel. Band intensity was normalized against the non-differential isoforms. Significant differences were calculated using a t-test (* p-value < 0.05).

**Fig. S5:**
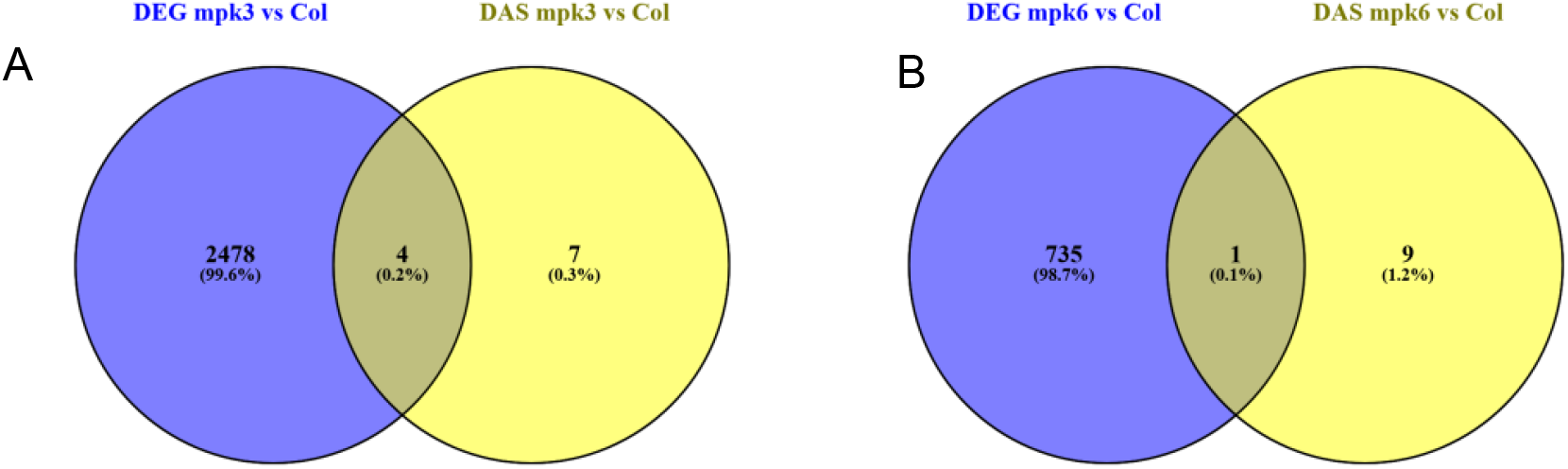
Venn comparison plot between differentially expressed and differentially spliced genes in *mpk3*, A or *mpk6*, B as compared to Col-0 wild type (FDR < 0.01)

